# A Comparison of Performance for Different SARS-Cov-2 Sequencing Protocols

**DOI:** 10.1101/2021.03.01.433428

**Authors:** Juanjo Bermúdez

**Affiliations:** Contignant Technologies SL c/Emigrant 30, Barcelona, Spain (08906)

**Keywords:** COVID-19, SARS-Cov-2, genome assembly, virus genome, genome sequencing, de novo genome assembly, sequencing protocols, ARTIC protocol

## Abstract

SARS-Cov-2 genome sequencing has been identified as a fundamental tool for fighting the COVID-19 pandemic. It is used, for example, for identifying new variants of the virus and for elaborating phylogenetic trees that help to trace the spread of the virus. In the present study we provide a comprehensive comparison between the quality of the assemblies obtained from different sequencing protocols. We demonstrate how some protocols actively promoted by different high-level administrations are inefficient and how less-used alternative protocols show a significant increased performance. This increase of performance could lead to cheaper sequencing protocols and therefore to a more convenient escalation of the sequencing efforts around the world.

## Introduction

There are two basic strategies to recreate a genome departing from the data obtained by the actually available sequencing machines:

1. Recreate the genome with no prior knowledge using de novo sequence assembly
2. Recreate the genome using prior knowledge with reference based alignment/mapping

It is generally accepted that each strategy has its own advantages and drawbacks. The quality of reference-based assembly is heavily dependent upon the choice of a close-enough reference: identification of some variantss can be missed if the sample is not close enough to the reference. In the other hand, de novo genome assembly is more computationally exigent and not always possible from the available data.

“Current variant discovery approaches often rely on an initial read mapping to the reference sequence. Their effectiveness is limited by the presence of gaps, potential misassemblies, regions of duplicates with a high-sequence similarity and regions of high-sequence divergence in the reference. Also, mapping-based approaches are less sensitive to large INDELs and complex variations” (1)

“We document that 18.6% of SNP genotype calls in HLA genes are incorrect and that allele frequencies are estimated with an error greater than ±0.1 at approximately 25% of the SNPs in HLA genes. We found a bias toward overestimation of reference allele frequency for the 1000G data, indicating mapping bias is an important cause of error in frequency estimation in this dataset.” (2)

“Detecting indels is challenging for several reasons: (1) reads overlapping the indel sequence are more difficult to map and may be aligned with multiple mismatches rather than with a gap; (2) irregularity in capture efficiency and nonuniform read distribution increase the number of false positives; (3) increased error rates makes their detection very difficult within microsatellites; and (4) localization, near identical repetitive sequences can create high rates of false positives” (3)

In an ideal scenario, researchers should have both options available: reference-mapping and de-novo assembly. If one of these is missed, the results do not count with the maximal possible reliability. And if there is the possibility to have both at the same cost, there is absolutely no reason for not having both.

For that reason it is important that the libraries for sequencing SARS-Cov-2 are designed with de novo genome assembly in mind.

Some studies have already been developed to assess the performance of the most commonly used protocols (4), but these are exclusively focused on the obtained coverage of the reads and not in the quality of the de novo assemblies. This study will establish a comparison of protocols based on the quality of the de novo assembly, which is a more exigent metric to asses the performance of the protocols. The performance of mapping to a reference genome will not be analyzed as this has already been analyzed in previous studies and a superior performance in de novo assembly is already strongly correlated to a superior performance in reference-mapping.

## Method

I used different search patterns at the NCBI SRA (5) website to find SARS-Cov-2 sequencing data obtained using different protocols. Despite this is not a totally reliable method (some search terms are ambiguous) I think it can help to understand the proportions.

Table 2 shows the number of matches found for every sequencing hardware technology. Despite some protocols were developed for some specific hardware, we can see how these are being used for other hardware too. for example, there are many more ARTIC (6) results for Illumina than for Nanopore despite the protocol was initially designed for Nanopore.

**Table 1.** Queries at the NCBI portal

| Protocol | Query |
| --- | --- |
| ARTIC | sars ARTIC |
| ARTIC V2 | sars ARTIC V2 |
| ARTIC V3 | sars ARTIC V3 |
| RANDOM | sars random NOT ARTICV3 NOT ARTICV2 NOT ARTIC |
| ALL | sars |

**Table 2.** Sequencing runs found for every protocol and hardware

| Protocol | Illumina | Nanopore | Capillary | LS454 | Ion Torrent | BGISEQ | PacBio |
| --- | --- | --- | --- | --- | --- | --- | --- |
| ARTIC | 78115 | 22909 | 12 | 4 | 0 | 0 | 0 |
| ARTIC V2 | 2681 | 66 | 0 | 0 | 0 | 0 | 0 |
| ARTIC V3 | 714 | 0 | 0 | 0 | 0 | 0 | 0 |
| RANDOM | 7525 | 323 | 0 | 1 | 26 | 109 | 0 |
| ALL SARS | 215187 | 34866 | 1762 | 7 | 536 | 148 | 25 |

See how results corresponding to the ARTIC protocol roughly correspond to 41% of all available SARS-Cov-2 runs in the SRA archive.

From the results for these queries I randomly selected some runs and downloaded the data sets. Then I assembled the data sets using the best performing genome assembly software from SPAdes (7), rnaSPAdes (8) and metaSPAdes (I will note as xSPAdes the best result obtained from these). In case the runs contained long reads Flye and Canu (9) was also applied. I finally assembled some of the short-read runs with Contignant s-aligner (10).

SPAdes, rnaSPAdes and metaSPAdes have been demonstrated to be the best-performing open-source software for viral genome de-novo assembly in different previous studies. Flye and canu are considered the best-performing assembly utilities for long-read data. Meanwhile, s-aligner is a new de novo genome assembler that has recently demonstrated superior performance for viral-genome assembly over the previous short-read assemblers.

## Results

Table 3 show the results obtained.

**Table 3.** Sequencing results for randomly selected data-sets

| Run Id | Hardware | Library design | layout | run size (MB) | Canu NG50 | xSPAdes NG50 | s-aligner NG50 |
| --- | --- | --- | --- | --- | --- | --- | --- |
| SRR12351628 | miseq | ARTIC V3 | paired | 84 |  | 4371 | 8431 |
| SRR12819233 | novaseq 6000 | random | paired | 382 |  | 21585 | 29845 |
| SRR12445029 | ion torrent | random | single | 1413 |  | 4980 | 29299 |
| SRR13684392 | miseq | ARTIC | paired | 1500 |  | 29404 |  |
| SRR11410529 | miseq | ARTIC | paired | 111 |  | 19294 |  |
| SRR12045777 | miseq | ARTIC v3 | paired | 188 |  | 19338 | 20522 |
| SRR13200927 | nextseq 500 | unspecified | single | 287 |  | 9412 | 10610 |
| SRR10903401 | miseq | random | paired | 140 |  | 4094 | 10610 |
| SRR12623307 | miseq | ARTIC v3 | paired | 11 |  | 19.283 |  |
| SRR11772204 | miseq | ARTIC v2 | paired | 113 |  | 29.837 |  |
| SRR12045770 | miseq | ARTIC v3 | paired | 100 |  | 1.412 | 19.242 |
| SRR11410528 | miseq | ARTIC | paired | 76 |  | 19.291 | 19.294 |
| SRR13660064 | miseq | ARTIC v3 | paired | 10 |  | 16.463 | 12.708 |
| ERR5094566 | gridion | liverpool | single | 37 | 0 | 4.216 |  |
| SRR13623050 | miseq | ARTIC v3 | paired | 143 |  | 29.842 | 29.814 |
| SRR13623049 | miseq | ARTIC v3 | paired | 131 |  | 29.833 |  |
| SRR12481157 | miseq | random | paired | 94 |  | 23.583 | 29.836 |
| ERR4182482 | GridION | unknown | single | 10 | 0 | 8.315 |  |
| SRR13574254 | illumina | ARTIC v3 | paired | 435 |  | 1.000 | 11.459 |
| SRR13727443 | illumina | artic v3 | paired | 850 |  | 1.631 |  |
| SRR13731834 | illumina | artic v3 | paired | 50 |  | 29.687 |  |
| SRR13495171 | illumina | random | paired | 650 |  |  | 29.820 |
| SRR13380666 | PacBio | hybrid | single | 78 | 0 |  |  |
| ERR5094578 | minion | artic v3 | single | 14 | 0 | 0 | 2.835 |
| ERR5165938 | nextseq 550 | hybrid | paired | 213 |  |  | 11.743 |
| SRR13380665 | PacBio | hybrid | single | 633 | 15.123 |  |  |
| ERR5165938 | nextseq 550 | hybrid | paired | 605 |  | 29.839 |  |
| SRR13727440 | illumina | ARTIC V3 | paired | 1126 |  | 0 | 12.591 |
| SRR12445036 | ion torrent | random | single | 191 |  | 5.104 | 29.112 |
| ERR4971211 | nextseq 500 | random | paired | 126 |  |  |  |
| SRR13615951 | BGISEQ | random | single | 48 |  | 29.858 | 29.846 |
| SRR13615945 | BGISEQ | random | single | 8 |  | 29.852 | 29.797 |
| SRR13615944 | BGISEQ | random | single | 3 |  | 29.852 | 29.829 |
| SRR13615947 | BGISEQ | random | single | 5 |  | 29.856 | 29.837 |
| SRR13615942 | BGISEQ | random | single | 40 |  | 28.307 | 29.754 |
| SRR13300938 | ion torrent | random | single | 586 |  |  | 18.500 |
| SRR12445040 | ion torrent | random | single | 405 |  | 5.560 | 29.340 |
| SRR12445032 | ion torrent | random | single | 119 |  | 2.475 | 29.351 |
| SRR13050769 | hiseq | random | paired | 3659 |  | 0 | 25.854 |
| SRR13495171 | illumina | random | paired | 651 |  | 0 | 29.804 |

Results in which both assembling methods under performed were excluded as likely due to problems in the data-set.
Empty cells correspond to assemblies that were not tried because of lack of relevance for the study.

**Table 4.**
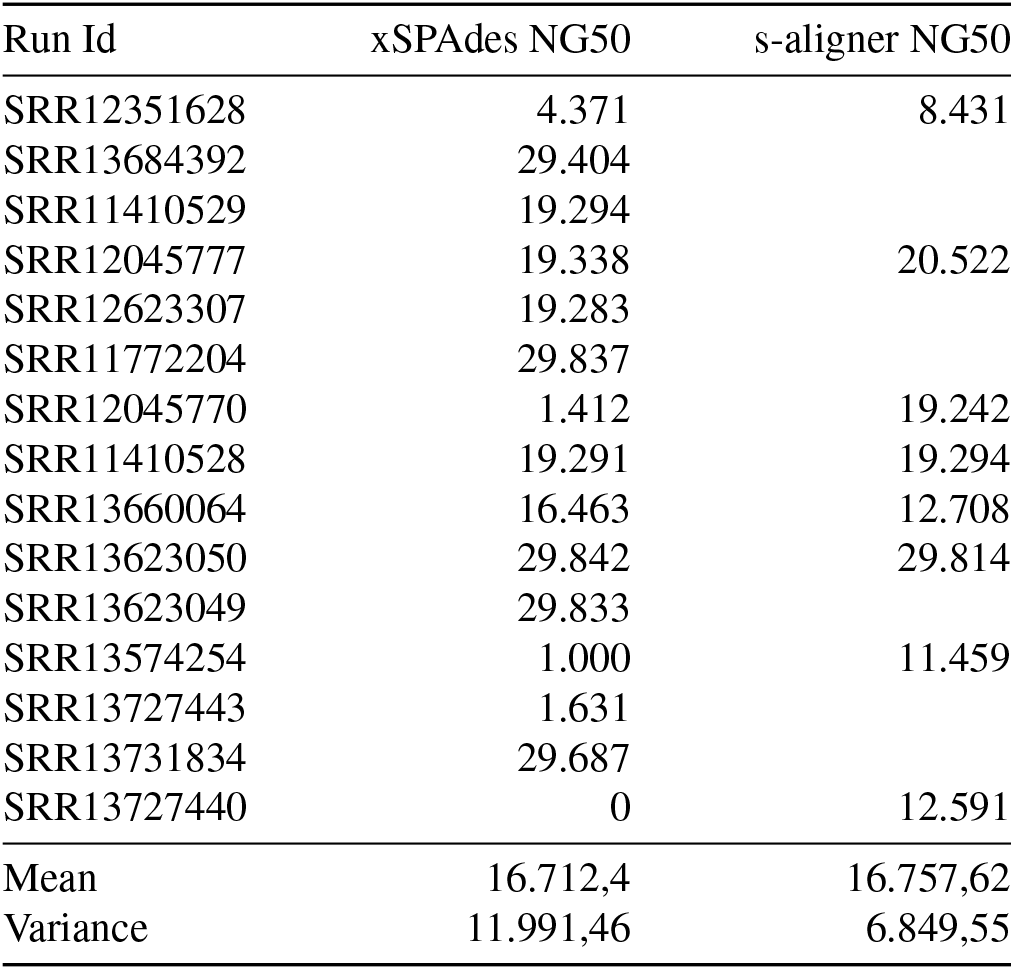
Sequencing results for runs obtained from the ARTIC protocol

**Table 5.** Sequencing results for runs obtained with random primers amplification

| Run Id | xSPAdes NG50 | s-aligner NG50 |
| --- | --- | --- |
| SRR12819233 | 21.585 | 29.845 |
| SRR12445029 | 4.980 | 29.299 |
| SRR10903401 | 29.877 |  |
| SRR12481157 | 23.583 | 29.836 |
| SRR12445036 | 5.104 | 29.112 |
| SRR13615951 | 29.858 | 29.846 |
| SRR13615945 | 29.852 | 29.797 |
| SRR13615944 | 29.852 | 29.829 |
| SRR13615947 | 29.856 | 29.837 |
| SRR13615942 | 28.307 | 29.754 |
| SRR13300938 |  | 18.500 |
| SRR12445040 | 5.560 | 29.340 |
| SRR12445032 | 2.674 | 29.351 |
| SRR13050769 | 0 | 25.854 |
| SRR13495171 | 0 | 29.804 |
| Mean | 17.220,57 | 28.654,93 |
| Variance | 13.063,15 | 2.984,97 |

From these results, some observations can be extracted.

### A. Short-read data-sets outperform long-read ones

I still have not found a long-read data-set that completes a perfect assembly. Doesn’t matter the library design or the technology employed (Nanopore or PacBio). The mean NG50 for long-read data-sets is 7.622 while any protocol using short-reads at least doubles that.

In addition, the obtained sequences have a higher misassembly rate, which makes that data less feasible for variant detection.

### B. The ARTIC protocol is far from delivering optimal results

Despite being widely used (41% of runs in the SRA archive) its performance is low and far from the best-performing protocols. If we only consider results for shortread data the mean NG50 is 16.712, which is a quite bad result.

### C. The ARTIC protocol doesn’t outperform other protocols

When making use exclusively of open-source assembly software, the ARTIC protocol doesn’t even significantly outperform results from other protocols. Its NG50 mean is similar to the NG50 overall mean of all protocols using opensource software: 16.712 with ARTIC vs 15.865 overall, and slightly lower than protocols using random primers (17.220).

### D. Library designs with random primers largely outperform designs with fixed primers when using s-aligner

When making use of all available software options, not only open-source, designs with random primer selection largely outperform designs with fixed primer selection, like ARTIC. If we compare the NG50 mean from results for shortread data employing ARTIC and SPAdes (16.712), it is a 71% lower than the NG50 mean obtained from random-primer data and s-aligner assembler (28.654). Indeed, the combination of s-aligner plus random-primer data guarantees in most cases an almost perfect assembly of the virus genome. Thirteen out of fifteen cases got as result an almost-perfect assembly.

This observation is corroborated by the frequent presence of gaps in the reference-mapping of runs obtained from fixed-primers designs. This is, indeed, something that could be expected from designs based on fixed primers. That limitation is already recognized by the WHO (11).

### E. The ARTIC protocol under-performs even when using s-aligner as software for genome assembly

S-aligner is, in general, a better tool for viral genome assembly. But even when using it, the ARTIC protocol underperforms compared to other protocols. The average NG50 using s-aligner for ARTIC data-sets is 16.757, which is similar to the average NG50 with open-source software (16.712), but far from the average NG50 obtained with s-aligner for random-primer protocols (28.654).

### F. No paired-read performance benefit over single-read

When using s-aligner as assembly software with random-primer library designs, there is no significant difference between using paired-end data or single-read data: 28.394 (single) vs 28.654 (overall).

## Conclusions

There are significant differences of performance between different protocols for sequencing the SARS-Cov-2 (figure 1). The difference of performance between using the ARTIC protocol with short-read technologies and using a random-primer design with s-aligner is statistically significant, with p-vakue <0.00001. The difference of performance in the NG50 metric is on average 71,5%. In addition, when evaluating the perfect-assembly ratio, we find that ARTIC has a 33,3% success rate, while the s-aligner-based protocol has a 86,7% success rate. With long-read data-sets, the success rate of ARTIC is 0% and NG50 can’t even be calculated because of lack of data.

**Fig. 1.**
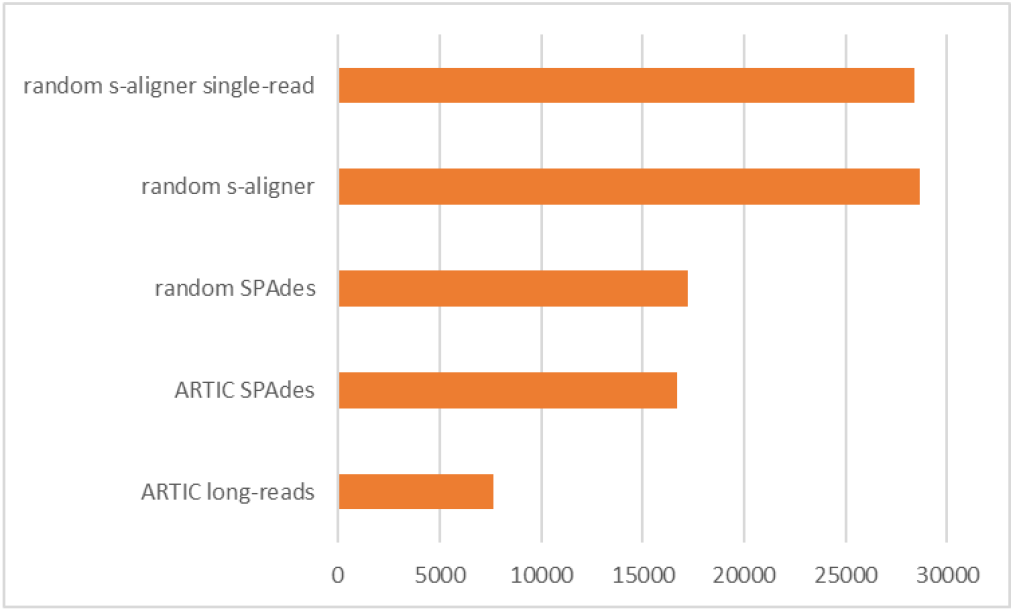
NG50 for different clusters of runs.

These results suggest that the hundreds of thousands of genome sequencing’s being done in the world to trace the spread of the virus and detect new variants are not making use of the most reliable and efficient methods. The low NG50 and perfect-assembly ratio suggest that these methods are even far from being reliable if de novo genome assembly is considered a need, as it is suggested by previous studies on the efficacy of only-mapping assembly. Mapping the data to a reference genome is usually considered a necessary but insufficient step, and it is always preferable to have a de novo assembly, being the only reason for not preferring that the unavailability of that possibility. We demonstrate in this study that there are protocols that reliably permit us to obtain de novo genome sequencing’s of SARS-Cov-2: a tool that would improve the quality of the actual efforts to trace the virus worldwide.

## Discussion

Another factor for considering which protocols to use for sequencing SARS-Cov-2 is the cost. ARTIC was specifically designed to be low-cost for that reason.

When evaluating the costs of different sequencing protocols three aspects should be considered.

1. The cost of the sequencing hardware
2. The cost of the products per sample
3. The overall time expended per sample

Unfortunately, I don’t have the necessary experience nor access to materials to evaluate these costs. For that reason I contacted several public-health organizations, warning them of the significant lack of performance of some protocols and offering them cooperation to find better ones. You can see on Annex I a list of entities that were contacted. None of them have acceded to cooperate at the moment of writing this manuscript. One can guess what their motivations are, but some motivations can be firmly discarded: they are not rejecting that because they are already developing equivalent studies nor because they already have the answers that such study would bring.

Even though I lack the experience to make a full analysis of the cost-effectiveness of different protocols for sequencing SARS-Cov-2, some clues can be extracted from the data in this study. We see how we can obtain reliable, almost-complete, de novo genome assemblies from data-sets under 10MB (therefore largely multiplexable), obtained with less-expensive hardware like Ion Torrent or BGI. Also with Illumina, we can establish cost-effective protocols making use of less data and single-read technology. That suggests that costeffective protocols are possible that are also reliable under a-de-novo assembly perspective and not only under a reference-mapping one. The increase of performance also suggests that a higher percentage of sequencing efforts will end-up in conclusive results, therefore eliminating the cost of most inconclusive results. All that information suggest that overall more cost-effective protocols than ARTIC are possible and desirable.

## Data availability statement

The data underlying this article are available as DOI: 10.5281/zenodo.4558343.

The s-aligner software is available for free at https://contignant.com for first-time users. It’s free to use for 15 days after installation. No personal identification is required but a contact email must actually be provided for downloading it.

## Supplementary Note A: Institutions invited to cooperate

See in Table 6 the list of public institutions that were contacted whether to warn them of a possible inefficiency in the applied protocols for sequencing SARS-Cov-2 (including an offer to cooperate) or to warn them of the existence of a new tool that could have an impact on the protocols for sequencing SARS-Cov-2 (offering them also cooperation).

**Table 6.** Public institutions contacted to warn them of a possible improvement in public protocols for the management of the COVID-19 crisis.

| Organization | City | Contact method | Contact date | Message content | Response |
| --- | --- | --- | --- | --- | --- |
| Hospital Universitari de la Vall d'Hebron | Barcelona (Spain) | Cold email | Feb 15, 2021 | Warning of low performance of ARTIC protocol. | They did not acknowledge reception |
| Hospital Clínic | Barcelona (Spain) | Cold email | Feb 15, 2021 | Warning of low performance of ARTIC protocol. | They did not acknowledge reception |
| Hospital Sant Pau | Barcelona (Spain) | Email to a connection and cold email to leaders | Feb 15, 2021 | Warning of low performance of ARTIC protocol. | Unofficially: not interested / not their scope. No official acknowledge of reception. |
| Sanger Institute | Hinxton (UK) | Cold email | Feb 15, 2021 | Warning of low performance of ARTIC protocol. | They acknowledged reception and opened a ticket. No further news from them. |
| Red Española de Investigación en Sida | Hinxton (UK) | e-mail recommended by a connection | Feb 9, 2021 | Warning of low performance of SARS-Cov-2 sequencing. | Rejected: too busy. |
| Barcelona Super-computer Center | Barcelona (Spain) | Cold email | Sept 2020 | Warning of better performance for viruses. | They did not acknowledge reception |
| Elixir Spain | Barcelona (Spain) | Cold email | Sept 2020 | Warning of better performance for viruses. | They did not acknowledge reception |
| Instituto Nacional de Biotecnología | Barcelona (Spain) | Cold email | Sept 2020 | Warning of better performance for viruses. | They did not acknowledge reception |

Several private institutions were also contacted. None has either responded.

